# G-LATO: Inference of Spatial Latent Ordering via Deep Gaussian Processes

**DOI:** 10.64898/2026.06.23.734031

**Authors:** Marcello Zago, Soham Mukherjee, Jan T. Schleicher, Paul-C. Bürkner, Ghazaleh Tabatabai, Manfred Claassen

## Abstract

Spatial transcriptomics enables the study of cells within their native tissue context, yet identifying gradients of cellular development remains challenging. We introduce a deep Gaussian process model to address this gap. Our method recovers spatially smooth gradients explaining observed gene expression. We illustrate our method on healthy liver and glioblastoma data in reconstructing known spatial organisation and uncovering new pathological gradients, thus providing robust inference for spatial biology.

## Main text

Biological processes unfold both in time and space, as exemplified by developmental programs such as organogenesis or pathological processes like immune infiltration in tumours. While spatial transcriptomics now enables the study of these spatio-temporal dynamics within their native tissue context, there is a lack of computational tools to learn spatial latent orderings, i.e., spatial cell state gradients, to describe such processes in detail. Spatial transcriptomics platforms are broadly categorised into sequencing- and imaging-based techniques, each with trade-offs between subcellular resolution and gene coverage [1].

Spatial latent ordering inference refers to extracting spatio-temporal dynamics from the data and closely mirrors pseudotime approaches for suspension-based single-cell RNA sequencing [2, 3]. However, current approaches are constrained by their reliance on an initial cluster assignment [4], the reconstruction of disjoint local gradients [5, 6], or the use of black-box architectures [7], thus hindering the interpretability of underlying biological systems. To overcome these limitations, we introduce **G-LATO** (**G**aussian process **lat**ent **o**rdering), a deep Gaussian process (GP) model [8] designed to infer a latent ordering in spatial transcriptomics data (Fig. 1a). This latent ordering or latent time (denoted by *t*) generalises pseudotime [3] to cellular neighbourhoods, explicitly modelling spatial continuity through a deep GP structure. The first layer of the deep GP model maps spatial coordinates to *t*, and the second layer maps *t* to gene expression dynamics via a multi-output GP. Together, these layers yield an interpretable model that produces spatially coherent spatial latent orderings of tissue dynamics.

**Fig. 1.**
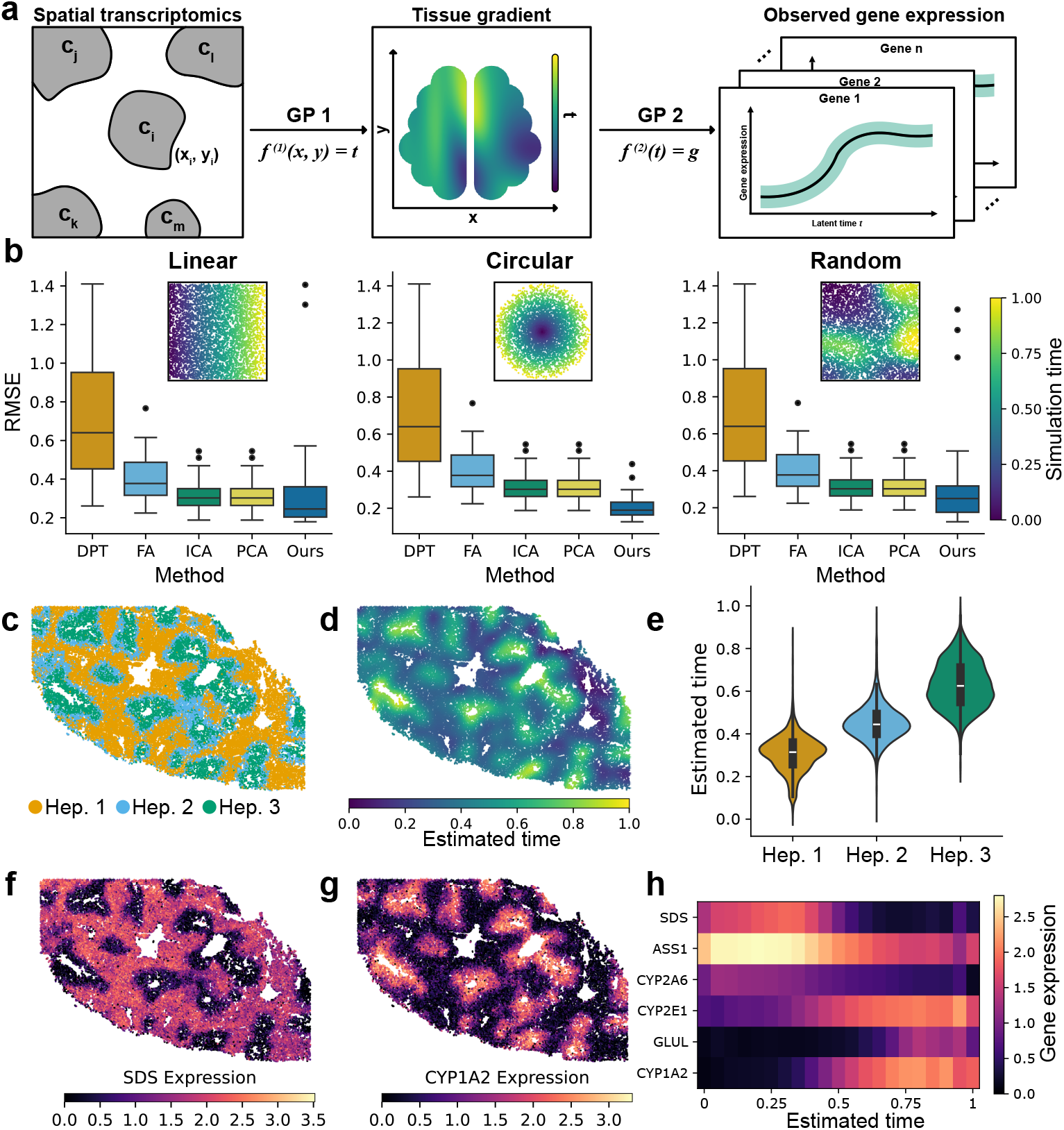
G-LATO consistently recovers smooth spatial latent orderings. **(a)** Schematic overview of the deep GP model. **(b)** Box-plots of root-mean-square error (RMSE) benchmark results for all methods. Inset scatterplots coloured by ground-truth simulation time show representative simulated spatial transcriptomics datasets. **(c, d, f, g)** Spatial distribution of hepatocyte subtypes, inferred latent time, *SDS* expression, and *CYP1A2* expression, respectively, across the tissue section. **(e)** Distributions of inferred latent time values across hepatocyte subtypes. **(h)** Heatmap of marker gene expression ordered by inferred latent time. Abbreviations: diffusion pseudotime (DPT), factor analysis (FA), independent component analysis (ICA), and principal component analysis (PCA).

We first assessed spatial latent order inference through a detailed simulation study featuring three distinct spatial patterns (Fig. 1b), and benchmarked G-LATO against various linear baselines and standard diffusion pseudotime analysis. Compared to these baseline methods, our model achieved higher linear recovery of the ground-truth simulation time, producing latent *t* estimates with higher accuracy (Fig. 1b) and strong association (Fig. B1). Concretely, we demonstrated that our proposed model reliably recovers latent ordering and outperforms the benchmark methods.

Next, we illustrate how our model recovers spatial latent orderings reflecting physiological gradients in real-world data settings. First, we considered a multiplexed FISH dataset of healthy liver tissue [9]. Hepatocytes in liver lobules are organised radially between portal triads and central veins, forming zonations with distinct metabolic activities (Fig. 1c) [9]. Motivated by this zonated organisation, we restricted our analysis to hepatocytes (*N* = 22, 912) and 95 highly expressed genes (see Online methods). G-LATO accurately reconstructed the repetitive lobular structure of the hepatocytes (Fig. 1d, e). Known zonation markers further validate the gradient, with peri-portal markers *SDS* and *ASS1* peaking at lower latent *t* values and peri-central markers *GLUL* and *CYP1A2* at higher values (Fig. 1f-h). This alignment between the marker expression and the latent time supports the interpretation that the inferred process faithfully encodes the functional metabolic zonation of hepatocytes.

In the second case study, we aimed to identify spatial latent ordering in a cancer context and considered a spot-based spatial transcriptomics dataset of glioblastoma featuring histologically annotated regions [10]. This dataset, comprising 3,213 spots and 152 highly expressed genes (as detailed in Online methods), encompasses distinct regions annotated as tumour core, transition zone, and infiltrative tissue (Fig. 2a). G-LATO reconstructed a smooth, continuous sequence of cell states with low values corresponding to infiltrative tissue and higher values to tumour core (Fig. 2b, c). Notably, the learned gradient in the transition zone appears as a continuous rather than a discrete pattern, indicative of gradual shifts in cell-type composition and metabolic activity across the tissue architecture (see Appendix D). Beyond the broader histological zones, the tumour core showed significant variability in latent time (Fig. 2b), suggesting the presence of transcriptionally distinct subregions. To investigate this heterogeneity, we partitioned the tumour region using a latent time threshold of *t* = 0.95 (Fig. 2b, red line) and performed a gene set enrichment analysis using the Hallmark gene sets [11]. The top-6 enriched pathways for each subregion (Fig. 2d) revealed that the low-latent-time region represents a highly proliferative tumour state with evidence of DNA damage. In contrast, the high-latent-time subregion appeared to represent a dying, immune-active niche characterised by immune signalling, apoptosis, and complement system pathways. These results show that G-LATO effectively scales to spot-based spatial transcriptomics data and provides a robust, exploratory method for identifying and characterising new tissue subdomains.

**Fig. 2.**
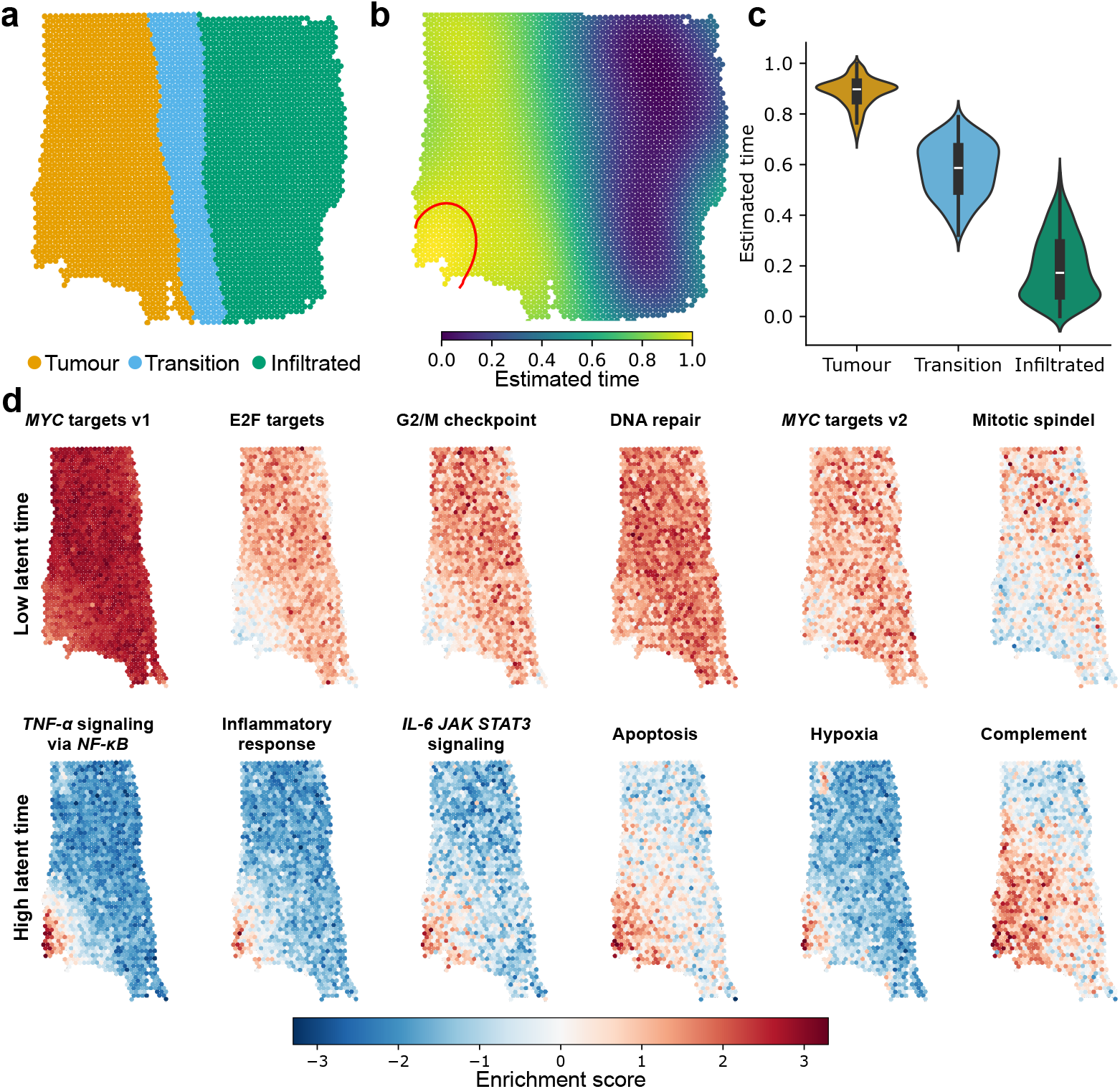
Recovering disease-associated latent orderings in glioblastoma reveals intratumour heterogeneity. **(a)** Spatial distribution of the histological annotations (tumour core, transition zone, infiltrative tissue) across the tissue section. **(b)** Inferred latent time from our method, revealing a global gradient of tumour infiltration. Binary splitting of the tumour core region for gene-set enrichment with threshold *t* = 0.95 is shown in red. **(c)** Distributions of inferred latent time values across histological region. **(d)** Spatial maps of enrichment scores for top Hallmark gene-sets in the tumour core. Top row: gene sets enriched in the low-*t* tumour subregion (*t <* 0.95), bottom row: gene sets enriched in the high-*t* subregion (*t >* 0.95).

In conclusion, G-LATO provides an interpretable and robust methodology for inferring spatially coherent latent cell state orderings across diverse biological contexts, including simulated data, liver zonation, and glioblastoma. The model demonstrates high versatility, performing effectively across both single-cell-resolved and spot-based spatial transcriptomics platforms, and successfully capturing diverse gradient architectures ranging from repetitive patterns to varying gradient scales. The primary strengths of our approach lie in ensuring spatially consistent gradients, inherent GP-based mechanistic interpretability, and computational scalability through variational inference (see Table C1). By eliminating the requirement for predefined root clusters of a dynamical process through previous biological knowledge, G-LATO enables the unbiased discovery of tissue gradients, making it a powerful tool for exploratory analyses of uncharacterised spatial domains. Although our case studies affirm the squared exponential covariance function as a robust default, yielding comparable results to alternatives like the Matérn 3/2 (see Appendix D), future work could investigate other covariance functions better suited for distinct biological patterns. Additional extensions to G-LATO include section-specific input GPs for joint analysis of multiple tissue sections, or adapting it to spatial proteomics data by modifying the output likelihood. Overall, our deep GP approach for spatial latent ordering inference has the potential for widely applicable integrated analysis of spatial multi-omics datasets.

## Online methods

### Gaussian processes

Gaussian processes (GPs) are a class of probabilistic methods for modelling unknown functions. GPs *f* (*x*) are specified by a mean function *µ*(*x*) and a covariance function *k* (*x, x*^*′*^) [12] for any arbitrary inputs *x, x*^*′*^ ∈ ℝ. We write *f* ∼ *GP* (*µ, k*) to indicate that *f* = *f* (*x*) is distributed as a GP such that any finite subset of *f* jointly follows a multivariate Gaussian distribution. Given an univariate output *y* and a single known covariate *x*, the standard GP regression is given by

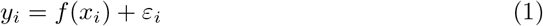

where (*y*_*i*_, *x*_*i*_) are paired samples for *i* = 1, 2, …, *N* with *N* being the sample size. *ε*_*i*_ is the *i*^*th*^ sample of the independent additive noise *ε* with error variance *σ*^2^ such that *ε* ∼ *N* (0, *σ*^2^). Thus, an univariate GP is written as

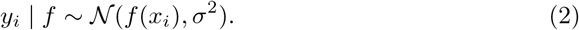

For a set of noisy observations *y*_1_, …, *y*_*n*_, the covariance between any two points *i* and *j* is defined as Cov(*y*_*i*_, *y*_*j*_) = *k*(*x*_*i*_, *x*_*j*_) + *σ*^2^*δ*_*ij*_, where *δ*_*ij*_ denotes the Kronecker delta. Thus, the covariance between distinct points is simply *k*(*x*_*i*_, *x*_*j*_), while the marginal variance for the *i*^*th*^ observation is Var(*y*_*i*_) = *k*(*x*_*i*_, *x*_*i*_) +*σ*^2^. Popular choices of *k* involve the Matérn class of stationary covariance functions [12]. Among them, we specifically consider the squared exponential (SE) and the Matérn 3/2 given by

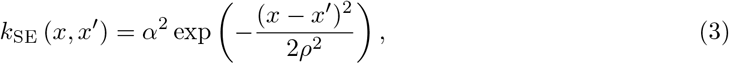

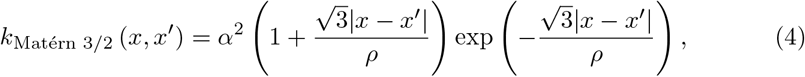

where *α* denotes the GP marginal SD, and the length scale *ρ* denotes the distance beyond which a pair of inputs can change significantly, thus determining the smoothness of functions *f* (*x*). For the main analyses of this paper, we use the SE covariance function, as it is a widely used choice [12]. Additional choices of covariance function involving the Matérn 3/2 for our model are highlighted in Appendix D.

In case of *P* -dimensional inputs (or *P* covariates) *x*_*i*1_, …, *x*_*ip*_ where *P* ≥ 2, the covariance functions are defined as

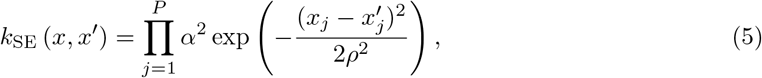

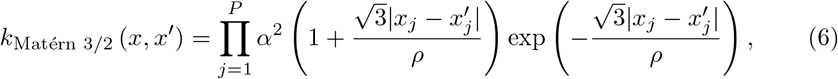

In our case, the covariance function parameters *ρ* and *α* are common across all the covariates. For stationary GPs, we use the Euclidean distance between observations for the *P* -dimensional covariates such that 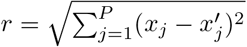. Then, a stationary covariance function is expressed as *k*(*r*). In the following section, we will use this stationarity property to approximate GPs.

### Hilbert space approximations

GPs are well known for their high computational complexity required to solve the Gram matrix *K* generated by the covariance function *k*. Specifically, for fitting a dataset with a sample size *N*, GPs have a computational complexity of *O*(*N* ^3^), which is prohibitive for large *N*. To circumvent this issue, we utilise the Hilbert space GP (HSGP) approximations [13, 14] where we approximate the covariance function *k* through its spectral decomposition computed from a finite set of representative basis functions.

The HSGP framework approximates a stationary covariance function *k*(*r*) over a bounded domain *x, x*^*′*^ ∈ Ω ⊂ [− *L*_1_, *L*_1_] × … × [− *L*_*P*_, *L*_*P*_] ⊂ ℝ ^*p*^ using a finite expansion of *M* ≪ *N* basis functions where *x, x*^*′*^ denotes *P* -dimensional covariates [13, 14]. For the SE and Matérn 3/2 with hyperparameters *α >* 0 and *ρ >* 0, the spectral densities are

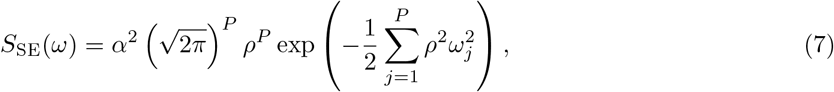

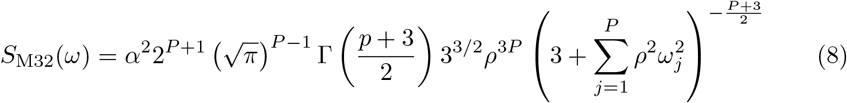

Based on the boundary conditions for *x, x*^*′*^ mentioned above, the number of basis functions *M* is decided such that 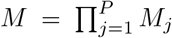 where *m*_*j*_ is the basis function for the *j*^*th*^ covariate. Then, a matrix S is constructed containing all the univariate combinations of *p*-tuples such that S ∈ *N*^*M ×P*^. For more details on S, see [14].

Using the combination of finite basis functions, the covariance function is thus approximated as

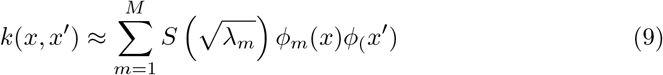

where the eigenvalues *λ*_*m*_ and eigenfunctions *ϕ*_*j*_(*x*) are given as

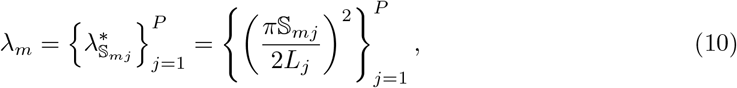

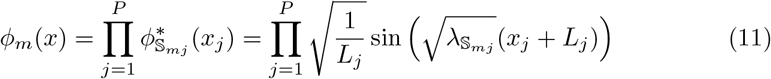

where *m* = 1, 2, …, *M*. The eigenfunctions and eigenvalues are obtained only based on the boundary conditions and are independent of the choice of covariance function. This decomposition allows the function *f* to be expressed as an additive model

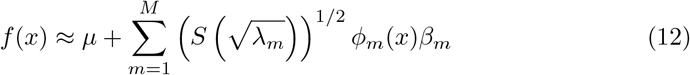

with *β*_*j*_ ∼ *N* (0, 1) [13]. Through the linear representation, the computational complexity is reduced from *O* (*N*^3^) to *O*(*NM*). For the multi-dimensional input setting of the HSGP approximation, we refer to Riutort-Mayol *et al*. [14]. For the HSGP conditions in the case of *P* = 1, see [14, 15].

### Multi-output HSGPs

Gene expression profiles can differ substantially across genes in expression amplitude, measurement noise, and dynamical change along a biological process. A single shared set of GP hyperparameters (*α, ρ, σ*) would thus not be able to model the underlying data sufficiently. Therefore, we model the *D* outputs jointly using a multi-output HSGP [15, 16], allowing each output dimension to learn its own hyperparameters.

Denoting by *y*_*di*_ the expression of gene *d* at observation *i*, we first specify an independent HSGP for each output *d* = 1, …, *D*,

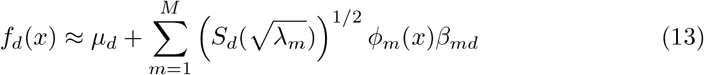

where *β*_*md*_ ∼ *N* (0, 1). To account for gene-gene (pairwise) covariance, the independent GPs are linked through a *D × D* across-output covariance matrix *C* [17, 18]. For each observation *i*, we obtain a vector of across-dimension correlated GP values as

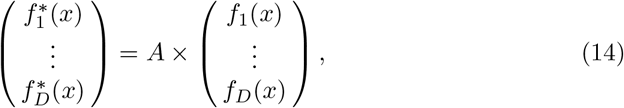

where *A* is the Cholesky factor of *C* [15, 16]. More information regarding the actual implementation can be found in Appendix A.

### The G-LATO model

The primary objective of this work is to recover a spatially consistent latent time *t* for cells or spots in spatial transcriptomics data. This requires introducing the latent *t* as input to a GP that models the observed gene expression. Previous latent variable extensions for HSGPs assume a prior-like structure for each latent variable derived from the experimental time of an scRNAseq experiment. [15, 16].

Since this temporal information is absent in spatial transcriptomics slides, and a spatial smoothness constraint is required, a deep GP model is more suitable [8].

Analogous to neural networks, deep GPs concatenate individual GPs, where the output of one layer serves as the input of the subsequent layer. We propose a two-layer model that leverages HSGPs to achieve significant computational gains across both layers.

The first layer models a function *f* ^(1)^ : (*c*_1_, *c*_2_) → *t* mapping two-dimensional spatial coordinates (*c*_1_, *c*_2_) onto the unidimensional latent time *t*. Utilising the SE covariance function imposes a smoothness prior on this mapping, ensuring high spatial continuity within the inferred latent ordering. The second layer models the function *f* ^(2)^ : *t* → *g*_*i*_, mapping the latent variable to observed gene expression for gene *i* to capture the dependency of transcriptional profiles along the biological process represented as the latent ordering. Following Mukherjee *et al*. [15], this layer is implemented as a correlated multi-output HSGP with dimension-specific parameters *α, ρ*, and *σ*.

To address the zero-inflation inherent in spatial transcriptomics data, our model uses a zero-inflated log-normal likelihood as yet another extension of the HSGP models shown in Mukherjee *et al*. [15]. By quantitatively modelling technical dropout from biological signal, this formulation enables the easy identification of robust genes that are less affected by strong dropout. Furthermore, a quantitative estimation of a gene’s zero-inflation can facilitate biological interpretation, allowing researchers to prioritise reliably expressed genes. In summary, these structural constraints of the model ensure the latent ordering *t* to capture continuous, spatially coherent cellular states, thereby enabling the reconstruction of biological trajectories, while remaining interpretable.

### Inference

The probabilistic model was implemented in Numpyro [19], enabling posterior inference of *t* and all GP parameters via stochastic variational inference (SVI) [20]. Given the observed gene expression *G* ∈ ℝ ^*N ×D*^ and a vector *θ*, representing the latent time and GP hyperparameters, we approximate the posterior *p* (*θ* | *G*) via SVI. This involves fitting a tractable guide *q*_*ψ*_(*θ*), where *ψ* represents the variational parameters. We utilised a mean-field normal approximation, which assumes a product of independent factorised Gaussians:

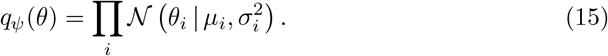

The variational parameters *ψ*_*i*_ = *{µ*_*i*_, *σ*^2^*}* are optimised by maximising the evidence lower bound

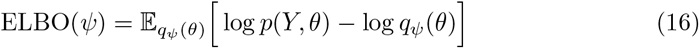

using Adam and a learning rate of 0.001 [21].

Although the posterior *p* (*θ* | *Y*) can also be estimated via Markov chain sampling algorithms, their high computational cost makes them unattractive for exploratory analysis of large-scale spatial transcriptomics data with cell and gene numbers up to the order of 10^4^ and 10^3^, respectively. The full generative model specifications and prior distributions are detailed in Appendix A. The final reported estimates of *t* were acquired by averaging over 1000 posterior draws, then scaling the result to the unit interval [0, 1].

### Simulation study

To assess model performance in recovering the latent time *t*, we conducted a simulation study with synthetic spatial transcriptomics data with known ground truth. We first generated 50 datasets using dyngen [22], a multi-modal simulator of single cells. All datasets consisted of 5000 cells and 500 genes undergoing a linear differentiation process. To mimic the spatial structures of real tissues, the simulated cells were projected into the two-dimensional space based on their simulation time, creating three distinct patterns (see Appendix B).

We benchmarked our proposed model against three classical latent variable methods (principal component analysis, independent component analysis, and factor analysis) [23] as well as diffusion pseudotime [3], a standard trajectory inference method for scRNA-seq data. Since the diffusion pseudotime algorithm depends on an initial state, we chose the cell with the lowest value in the first diffusion component. Since these benchmark models do not consider spatial information, we used only the observed gene expression as their respective input.

To assess the goodness of latent ordering recovery, we use the root mean square error (RMSE) of the normalised simulation time and normalised model estimates. Furthermore, for the quantitative evaluation of the linear recovery, we report the absolute Pearson correlation coefficient (see Appendix B).

### Real world case studies

To demonstrate the applicability and performance of our model on real-world data, we performed case studies on two distinct spatial transcriptomics datasets, one with single-cell resolution and one spot-based.

### Healthy Liver

For the single-cell resolution dataset, we analysed the spatial transcriptomics data from a healthy human liver [9], specifically sample AM042. Our analysis focused on recovering the functional liver lobule gradients defined by metabolically distinct hepatocyte populations. To isolate the biological signal, we subsetted the dataset to include only hepatocyte clusters. Spatially disconnected cells were removed by filtering out the top 1 % of cells with the highest distance to their nearest neighbour, resulting in a final dataset of 22,912 cells. The feature space was restricted to highly expressed genes detected in at least 50 % of cells (*D* = 95).

To account for the fine-scale structure of liver lobules relative to the overall tissue dimensions, we adjusted the prior distribution of the first GP length scale. Specifically, the mean of the length scale prior was set to 10 % of the mean pairwise distance between all cells, encouraging the model to capture the local spatial variation.

### Glioblastoma

Next, for the spot-based spatial transcriptomics dataset, we analysed a glioblastoma dataset [10], with the goal of recovering the pathological tumour border gradient. We explicitly analysed sample UKF269T, which contains three histologically distinct regions: tumour core, transition zone, and infiltrative tissue. To ensure a robust signal quality, the feature space was subsetted to retain only genes expressed in at least 75 % of the captured spots. The final dataset consisted of 3,213 spots and 152 genes. Given the lack of prior knowledge regarding the spatial scale of the gradients, we utilised the vague prior distributions for all model parameters (see Appendix A).

### Gene-set enrichment analysis

We performed a gene-set enrichment analysis using the Decoupler package [24]. Specifically, we used the univariate-linear model as the primary enrichment model. This approach was used to score the Hallmark gene sets defined by the MSigDB [11], allowing us to robustly estimate pathway activities based on the observed gene expression changes between the low-*t* and high-*t* tumour subregions of the glioblastoma dataset.

## Supplementary information

Supplementary material for the mathematical definition of the model can be found in the Appendix.

## Acknowledgements

M.Z. was supported by the Else Kröner Fresenius Stiftung (ClinBrAIn). S.M. was supported by DFG EXC 2180. J.T.S. was supported by DFG CL 792/1-1 and DFG EXC 2180. The authors thank the International Max Planck Research School for Intelligent Systems (IMPRS-IS) for supporting M.Z, S.M, and J.T.S.

## Declarations

- Funding: DFG CL 792/1-1, DFG EXC 2180 and Else Kröner Fresenius Stiftung (ClinBrAIn).
- Competing interests: M.C. is advisor to Vicinity Bio GmbH.
- Ethics approval and consent to participate: Not applicable
- Consent for publication: Not applicable
- Data availability: The public single-cell spatial transcriptomics dataset of healthy liver was obtained from datadryad under the doi: 10.5061/dryad.37pvmcvsg (https://datadryad.org/dataset/doi:10.5061/dryad.37pvmcvsg). The Visium dataset of Glioblastoma was acquired from the SPATA2 R-package with downloadSpataObject(sample name = “UKF269T”) [10].
- Materials availability: Not applicable
- Code availability: Model implementation and study details can be found here: https://github.com/MarcelloZago/G-LATO
- Author contribution:

- **Marcello Zago:** Conceptualisation, Formal Analysis, Methodology, Software, Visualisation, Writing - original draft, Writing - review & editing
- **Soham Mukherjee:** Conceptualisation, Methodology, Software, Writing - review & editing
- **Jan T. Schleicher:** Conceptualisation, Methodology, Software, Writing - review & editing
- **Paul-C. Bürkner:** Validation, Writing - review & editing
- **Ghazaleh Tabatabai:** Funding Acquisition, Supervision, Writing - review & editing
- **Manfred Claassen:** Conceptualisation, Funding Acquisition, Supervision, Writing - review & editing

## Appendix A

Full model specification

## Hyperparameters

The Hilbert space Gaussian process (HSGP) approximation requires two main user settings: the boundary condition *L* and the number of basis functions *m* [13, 14]. Following the recommendations of Riutort-Mayol *et al*. [14], for the first GP we computed *L*_1_ = *c · S* with *S* being the half-range of the centred data and the scaling factor *c* = 1.25. Since the spatial coordinates *c*_1_, *c*_2_ are bounded to [−0.5, 0.5] we get 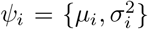. The inputs to the second GP layer follow a standard normal distribution and thus are technically not bounded. To account for this, we generously set *L*^(2)^ = 6, which covers 1 − 1.973 · 10^−9^ of the probability mass. Following the recommendations of Mukherjee *et al*. [15], we determine the number of basis functions as follows:

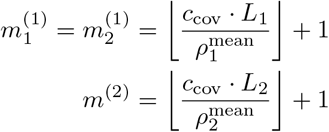

with the covariance scaling factor *c*_cov_ being 1.75 for the SE and 3.42 for the Matérn 3/2. Furthermore, *ρ*^mean^ denotes the mean of the length scale prior distribution of the respective GP layer. Additionally, we ensured the minimal number of basis functions for the first GP to be at least 5.

## Priors for the first GP layer

To assist the identifiability of the deep GP model, we omit noise from the first GP layer *f* ^(1)^ and fix its marginal standard deviation to *α*_1_ = 1. Consequently, only the length scale *ρ*_1_ is inferred, which determines the overall scale of the spatial gradients. Let *µ*_dist_ denote the mean Euclidean distance across all pairs of spatial input coordinates. To set the expected spatial correlation range to the tissue’s macroscopic scale, we centre the length scale prior at *ρ*_1_ ∼ *N*^+^(*µ*_dist_, 0.05), with ^+^ being the positive truncated normal distribution. If the expected spatial scale of the biological gradients disagrees with this macroscopic assumption, it is recommended to scale *µ*_dist_ accordingly, as demonstrated in the case study of the healthy liver tissue. As suggested in previous literature about the HSGP approximations, the weights for the basis functions have a standard normal prior distribution: *β*_1_ ∼ *N* (0, 1) [14, 15].

## Priors for the second GP layer

The second GP layer is a correlated multi-output GP with dimension-specific parameters *α*_2,*d*_, *ρ*_2,*d*_, and *σ*_2,*d*_ for *d* = 1, …, *D* genes, following the implementation from Mukherjee *et al*. [15]. The gene-specific parameters have the following prior distributions

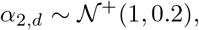

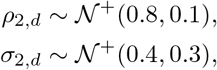

assuming the gene expression is normalised with the common log(*x* + 1)-transformation. For the Cholesky decomposition of the gene-gene correlation matrix *A*, we used the uniform LKJ distribution with a concentration parameter *η* = 1.0: *A* ∼ LKJCholesky_*D*_(1.0).

## Priors for the observational model

In addition to the previously defined gene-specific noise standard deviation *σ*_2,*d*_, the zero-inflated log-normal likelihood incorporates gene-specific intercepts *µ*_*d*_ and drop-out logits logit(*π*_*d*_). For log(*x* + 1)-normalised expression data, we assigned the following prior distributions across *d* = 1, …, *D* genes:

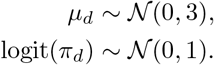

## Appendix B

Spatial mappings for the simulation study

This section details the procedure for generating the spatial gradients in the simulated spatial transcriptomics data. This process involves assigning a 2D coordinate (*c*_1_, *c*_2_)_*i*_ = (*x*_*i*_, *y*_*i*_) to each simulated cell *i* = 1, …, *N* based on its respective simulation time 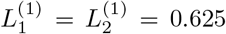. Because the previously defined hyperparameters and prior distributions (see Appendix A) require *x*_*i*_, *y*_*i*_ ∈ [− 0.5, 0.5], the following algorithms will adhere to this constraint. The linear and circular spatial patterns are generated using deterministic mapping functions with randomly sampled inputs as seen in Algorithms 1 and 2. For the random non-linear gradient, cells are assigned to coordinates on a random, smooth, Gaussian-filtered spatial gradient based on their simulation time 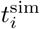 (Algorithm 3).

### Algorithm 1

Calculate linear spatial pattern

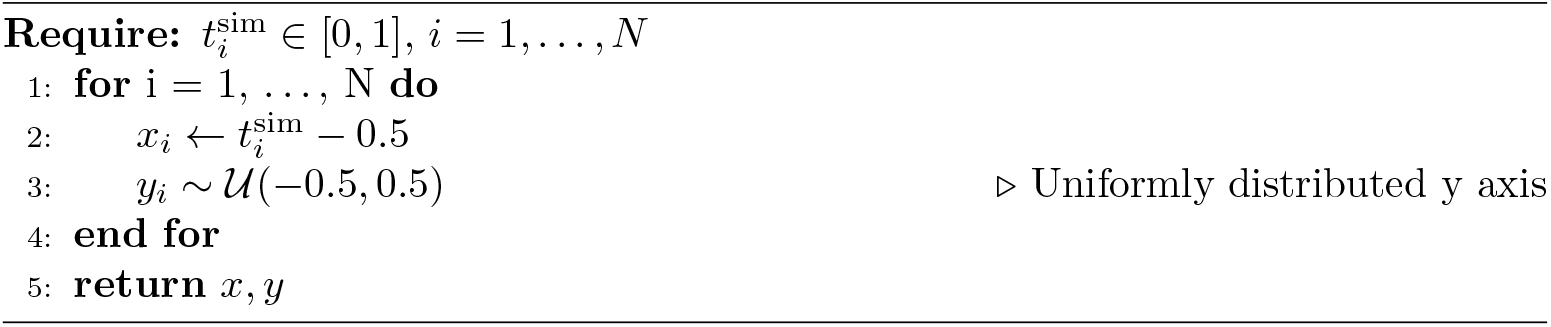

### Algorithm 2

Calculate circular spatial pattern

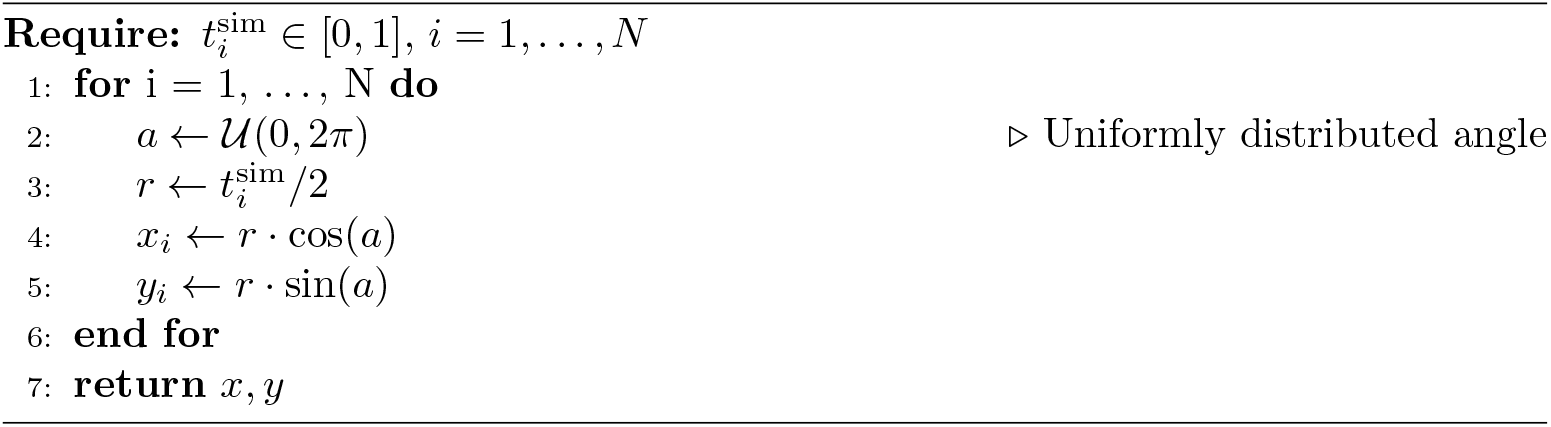

### Algorithm 3

Calculate random non-linear spatial pattern

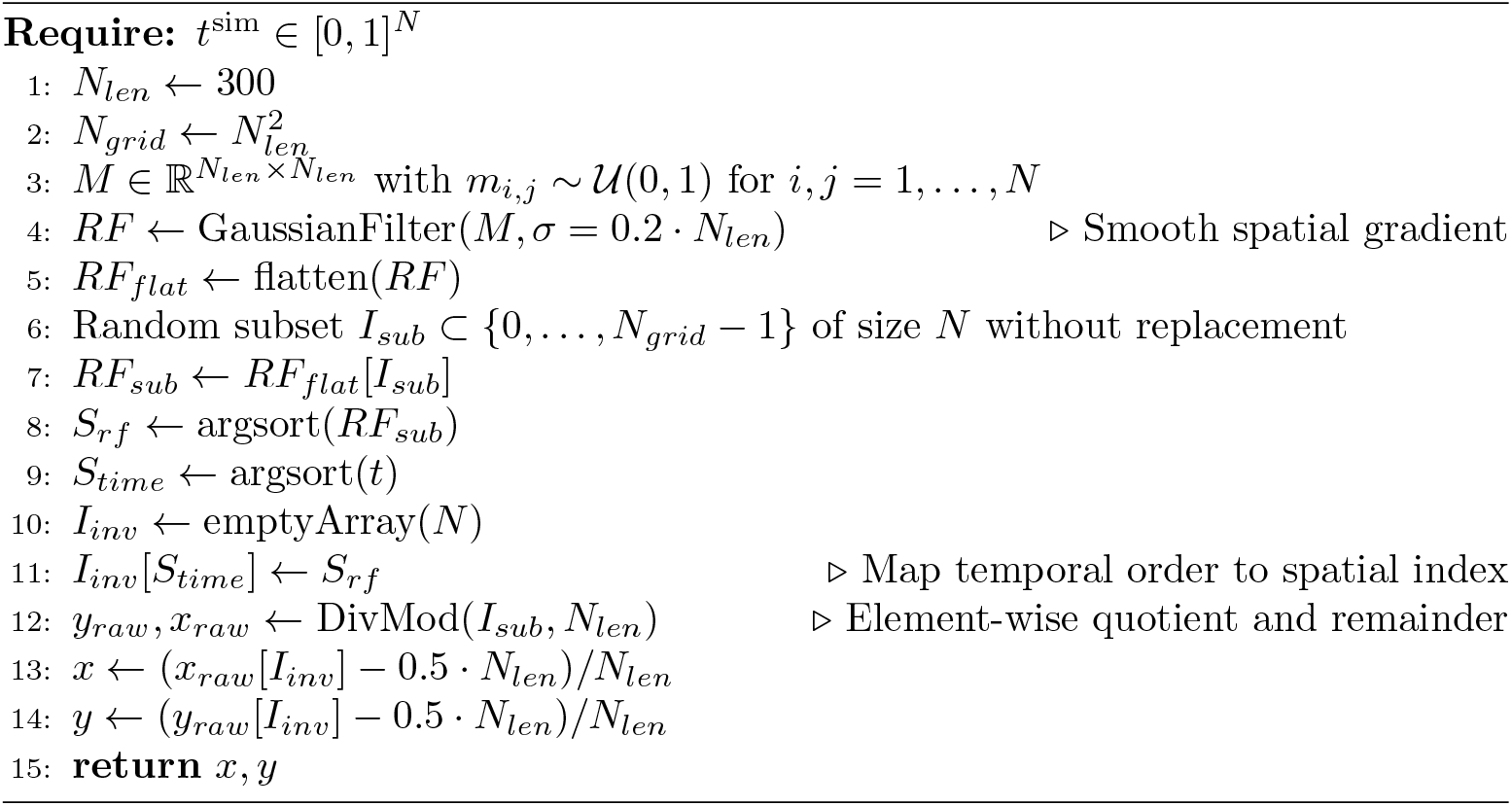

**Fig. B1.**
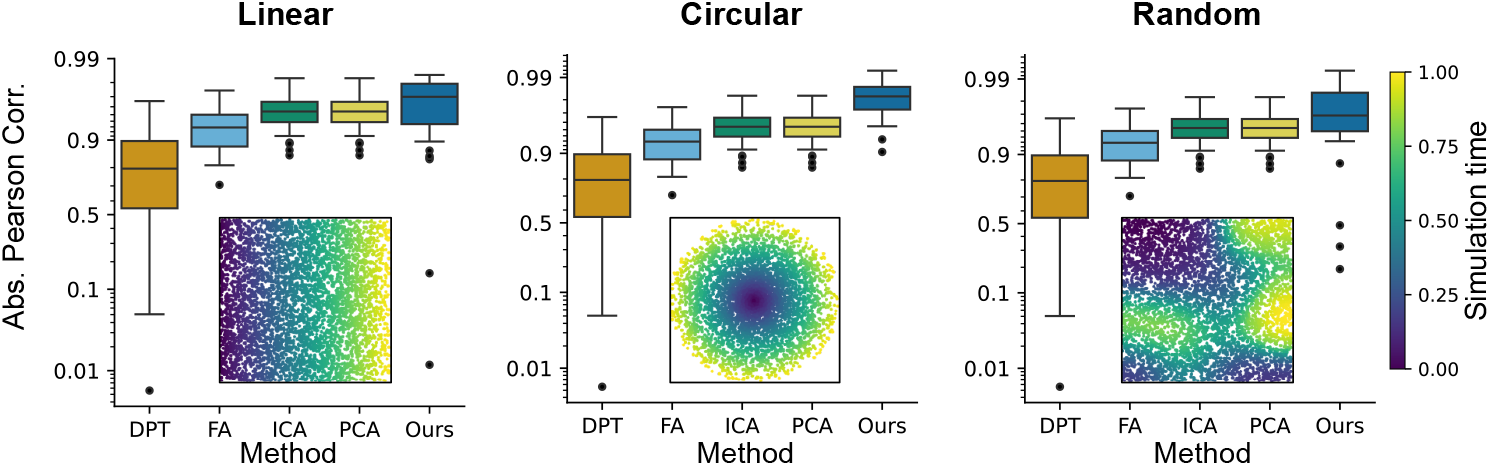
Simulation study absolute Pearson correlation. Box plots of the absolute Pearson correlation benchmark results for all methods. The y-axis is logit-scaled. Inset scatterplots coloured by ground-truth simulation time show representative simulated spatial transcriptomics datasets. Abbreviations: diffusion pseudotime (DPT), factor analysis (FA), independent component analysis (ICA), and principal component analysis (PCA).

## Appendix C

Runtime of inference

**Table C1.** Runtime performance across simulated and biological datasets.

| Dataset | # cells | # genes | Runtime (s) |
| --- | --- | --- | --- |
| Simulation (linear)* | 5000 | 500 | 215.8 ± 4.4 |
| Simulation (circular)* | 5000 | 500 | 216.5 ± 5.6 |
| Simulation (random)* | 5000 | 500 | 215.7 ± 4.5 |
| Glioblastoma | 3213 | 152 | 72.7 |
| Healthy liver | 22912 | 95 | 465.5 |

\*Results for simulation datasets are mean ± SD calculated over 50 iterations.

## Appendix D

Matérn 3/2 covariance function for spatial mapping

To assess the robustness of the proposed deep GP model with respect to the choice of covariance function of the input GP, we re-evaluated the case studies, substituting the squared exponential (SE) covariance function with the Matérn 3/2. The Matérn 3/2 covariance function is defined as

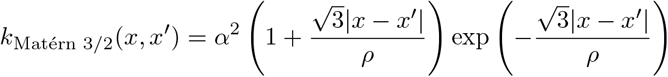

and has the following spectral density for input dimensionality *D* [14]:

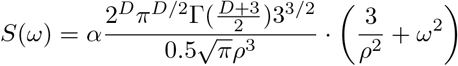

It induces a prior distribution over functions that are exactly once mean-square differentiable and thus less smooth [12]. In practice, the use of the Matérn 3/2 covariance function will introduce numerical instabilities during inference, especially for low length scales. To be able to perform an inference pipeline similar to that for the SE covariance function, it was necessary to reduce the initial value of the scale parameter of the variational guide to 0.001. For the glioblastoma dataset, this modification was sufficient to achieve stable evidence lower bound (ELBO) convergence. The resulting gradient (see Fig. D2) is qualitatively consistent with that obtained with the SE covariance function, resulting in a high-latent-time tissue corner (bottom-left) and a smooth gradient in the tumour core transition zone.

**Fig. D2.**
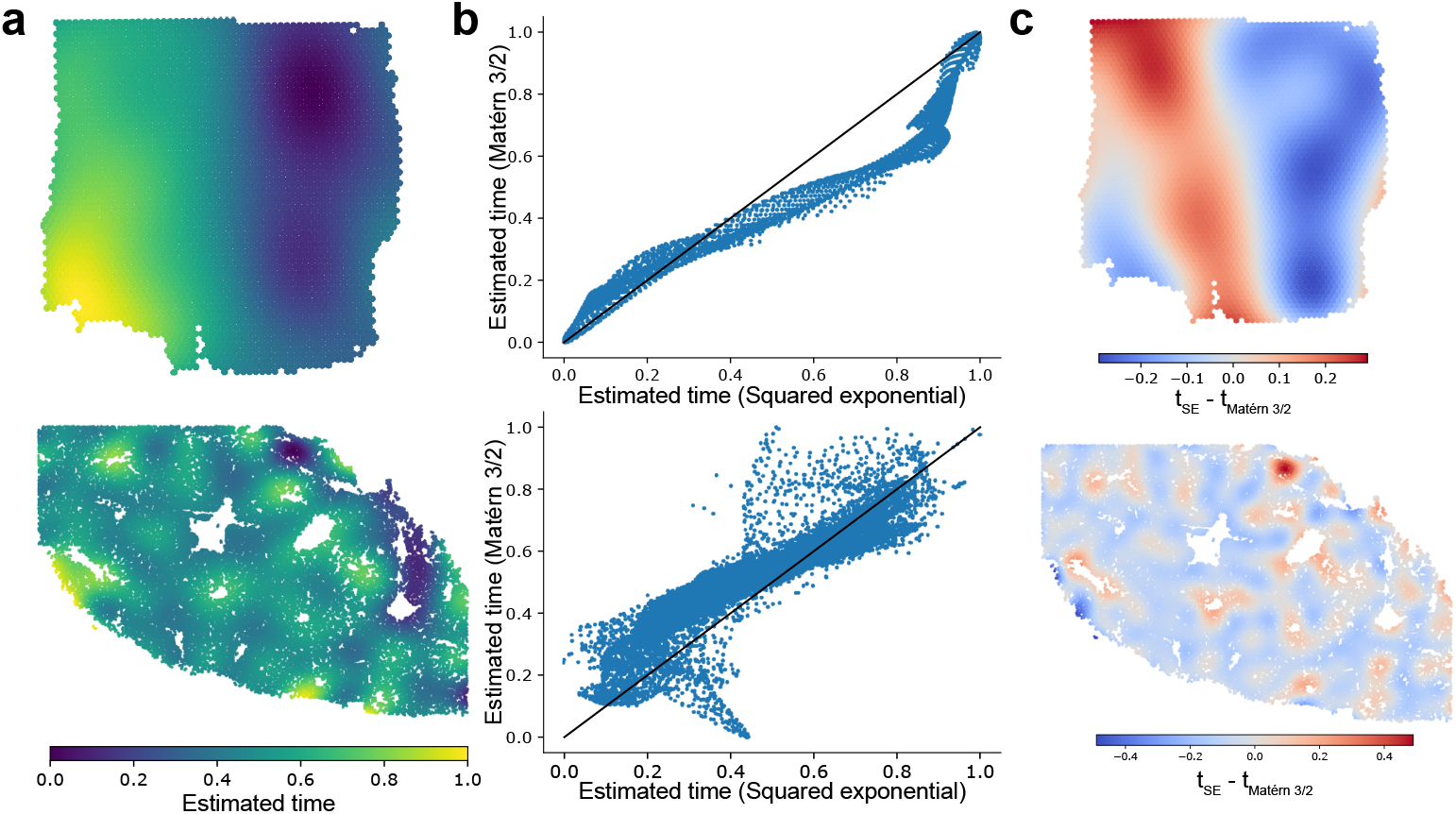
Matérn 3/2 input GP in case studies. **(a)** Inferred latent time for the Glioblastoma (top) and healthy liver (bottom) dataset over the respective tissue slides. Even with a less smooth covariance function, the transition zone of the glioblastoma dataset appears as a smooth rather than a discrete step-like gradient. **(b)** Scatter plots highlighting differences between estimates produced by the SE (x-axis) and Matérn 3/2 (y-axis) input GPs for the glioblastoma (top) and healthy liver dataset (bottom). **(c)** Spatial plots of the difference of inferred latent orderings (*t*_SE_ − *t*_Matérn 3/2_).

Model inference for the healthy liver dataset, which contains smaller gradients, proved considerably more challenging. Due to the challenging posterior geometry and numerical stability of the parameter space, the inference frequently resulted in a degenerate or undefined ELBO loss. To resolve this issue here, we increased the mean of the length scale prior distribution to one-third of the mean pairwise distance between all cells. The final resulting gradient map (see Fig. D2) captures the general trend of the liver lobules.

Overall, these findings suggest that the smoothness of the SE covariance function plays an important regularising role in the inference of tissue gradients. We therefore advocate retaining the SE as the default covariance function for the first GP layer and caution against the use of less smooth kernels unless numerical stability and prior calibration can be thoroughly validated on the dataset at hand.

